# One-Size-Fits-Many: Antisense oligonucleotides for rescuing splicing mutations in hotspot exons

**DOI:** 10.1101/2024.12.07.627366

**Authors:** Chaorui Duan, Stephen Rong, Luke Buerer, Christopher R. Neil, Juliann M. Savatt, Natasha T. Strande, William G. Fairbrother

## Abstract

Mutations that impact splicing play a significant role in disease etiology but are not fully understood. To characterize the impact of exonic variants on splicing in 71 clinically-actionable disease genes in asymptomatic people, we analyzed 32,112 exonic mutations from ClinVar and Geisinger MyCode using a minigene reporter assay. We identify 1,733 splice-disrupting mutations, of which the most extreme 1-2% of variants are likely to be deleterious. We report that these variants are not distributed evenly across exons but are mostly concentrated in the ∼8% of exons that are most susceptible to splicing mutations (i.e. hotspot exons). We demonstrate that splicing defects in these exons can be reverted by ASOs targeting the splice sites of either their upstream or downstream flanking exons. This finding supports the feasibility of developing single therapeutic ASOs that could revert all splice-altering variants localized to a particular exon.

## INTRODUCTION

RNA splicing is a critical step in the processing of eukaryotic genes, requiring multiple cis-elements for splice site recognition. These cis-elements encompass not only the 5’ and 3’ splice sites, branch points, and polypyrimidine tracts^1,2^ but also include splicing enhancers and silencers located within exons or introns^3–7^. They constitute a large mutational target for disease causing mutations^8^. Exonic mutations classified as synonymous or missense can also alter splicing by disrupting splicing enhancers or creating repressor sequences. It is estimated that a significant proportion, up to 60%, of human disease-causing mutations impact splicing rather than only affecting coding sequences^9^. Moreover, analysis of over 8,000 patients across 32 cancer types revealed that tumors exhibit up to 30% more alternative splicing events than normal samples. Additionally, thousands of splicing variants were absent in normal tissues^10^. This suggests that RNA splicing plays an important role not only in human diseases but also in cancer progression and metastasis. While extensive research has been conducted on identifying alternative splicing events as biomarkers for various cancers^11–15^, comparatively less focus has been placed on identifying splice-disrupting mutations, particularly within clinically actionable genes.

Antisense oligonucleotides (ASOs) have emerged as versatile therapeutic tools that can be designed to precisely target and modify RNA transcripts. ASOs are short oligonucleotides which offer a diverse range of mechanisms for treating genetic disorders based on their binding to specific RNA loci via Watson-Crick base pairing^16–18^. The first approved ASO therapy was introduced in 1998^19^. To date, 15 ASO drugs have received marketing approval from the United States Food and Drug Administration (FDA) for the treatment of human diseases^20^, including Duchenne muscular dystrophy (DMD)^21–23^, spinal muscular atrophy (SMA)^24–26^, amyotrophic lateral sclerosis (ALS)^27^, and other neurodevelopmental and neuromuscular disorders^28^. Although ASOs designed for allele-specific therapy have shown promise, identifying mutations and developing ASOs for each individual mutation can be time-consuming and costly. Here, we demonstrated that a single ASO can correct exon skipping caused by multiple mutations within the same exon, offering a versatile ‘one-size-fits-many’solution. This ASO strategy has the potential to be more cost-effective and represents a new frontier in therapeutic development.

In this study, we employed the Massively Parallel Splicing Assay (MaPSy) to evaluate exonic mutations in 71 clinically actionable genes, identifying 1,733 splice-disrupting variants. Furthermore, our findings revealed a non-uniform distribution of these splice-disrupting variants across exons. This observation highlighted the presence of hotspot exons that are particularly susceptible to such mutations. Importantly, the multitude of splicing mutations occurring in these hotspot exons can be efficiently rescued by a single exon-targeting ASO.

## RESULTS

### MaPSy detects splice-disrupting variants in clinically actionable genes

To assess the effects of reported variants on the splicing outcomes of exons within 76 clinically actionable, disease-associated genes^29^, we identified 36,471 single-nucleotide variants within the exonic regions of 71 genes from Geisinger MyCode and ClinVar. These regions were suitable for analysis using our Massively Parallel Splicing Assay (MaPSy), which was originally developed from an *in vitro* SELEX procedure designed to identify splicing enhancer motifs^30^. Due to oligonucleotide synthesis limitations, candidate variants could occur 1) in exons at most 120 nt in length and be incorporated into a full-length exon construct or 2) within 90 nt of an exon’s 3’ splice site and be incorporated into a half-exon construct containing a common 3’ segment and 5’ splice site (Figure 1a). Variants were synthesized using Agilent pooled oligo synthesis with the variant and wildtype species for each exon engineered into a three-exon, two-intron minigene reporter to construct the MaPSy library.

**Figure 1.**
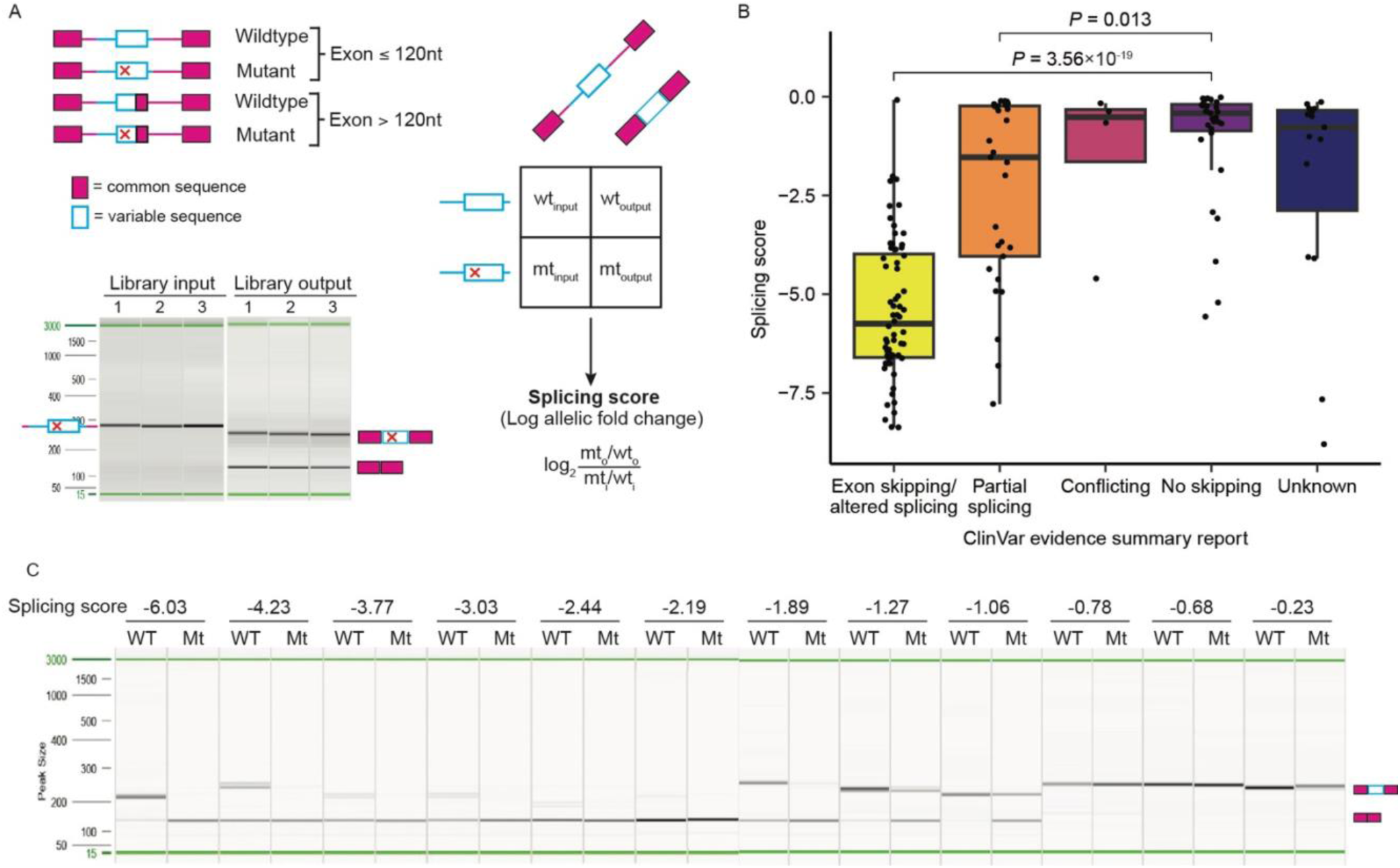
Massively parallel splicing assay identifies splice-disrupting variants in clinically-actionable genes. A) Exons containing variants from MyCode and ClinVar were engineered into splicing reporter minigenes alongside wildtype exons. Due to oligonucleotide synthesis limitations, exons longer than 120 nt were truncated and inserted into minigenes with a common 5’ splice site sequence. After transfection of the constructed input library into HEK293T cells, RNA was extracted and amplified with primers in the common upstream and downstream exons. Read counts from RNA-seq of input and output samples mapping to wildtype and mutant species allow for the calculation of allelic fold change for each variant. B) ClinVar evidence summaries for the tested variants were manually reviewed for reports that variants were associated with exon skipping/altered splicing, partial splicing or no skipping. Conflicting reports or those indicating an unknown effect on splicing are included for comparison. C) 12 variants with a range of allelic fold changes (listed above gel) were selected for PCR validation.

The input library was sequenced, and following transfection in triplicate into HEK293T cells, the library output was sequenced. The most common consequence of disrupted splicing is exon skipping (Figure 1a, gel). Comparison of input and output read counts for each species allows for the calculation of allelic fold change (mutant/wildtype score) (Figure 1a, right). We refer to the log_2_-transformed allelic fold change as the MaPSy splicing score, where a negative score indicates that the variant splices worse than the wildtype sequence. Read counts for the MaPSy experiment were highly correlated between input and output replicates (R > 0.99, Supplementary Fig. 1).

For the assayed variants, ClinVar data was manually reviewed for reports that variants had effects on splicing. Variants which were reported to cause exon skipping/altered splicing or partial splicing had more negative MaPSy splicing scores than variants which were reported to not cause skipping, confirming that MaPSy results are predictive of exon skipping in endogenous contexts (t-test, *P* = 3.56×10^-19^ for exon skipping, *P* = 0.013 for partial splicing; Figure 1b). Additionally, we selected 12 variants with a range of MaPSy splicing scores for cloning and validation with RT-PCR (Figure 1c). Variants with highly negative MaPSy splicing scores (<-2) were associated with complete exon skipping events, while less disruptive variants resulted in a mixture of skipping and inclusion. These results suggest that MaPSy is robust at detecting splice-disrupting variants (SDVs), particularly those associated with exon skipping.

### Splice-disrupting variants are associated with pathogenicity and exhibit evidence of purifying selection

Due to the sensitivity of minigenes relative to endogenous gene processing, our analyses of MaPSy results have focused on variants which exceed a critical splicing score threshold^31,32^. Defining splice-disrupting variants as those with a MaPSy splicing score below −1.5 at a 5% false discovery rate, 1,733 variants out of the 32,112 variants which passed quality checks (5.4%) were identified as SDVs. However, as seen in the variant validation presented in Figure 1c, some variants close to this threshold may not be clinically relevant as they only induce exon skipping to a limited extent. In order to focus on the variants that are most likely to cause severe splicing defects, we also conducted analyses on the variants with MaPSy splicing scores in the bottom 2% of the distribution of all variants passing quality checks. We found these variants, which we term extreme splice-disrupting variants (eSDVs), to be associated with stronger predicted splicing defects and higher levels of conservation (Supplementary Figure 2).

We found eSDVs to be categorized as pathogenic or likely pathogenic in ClinVar at a higher rate than other variants (Figure 2a). Analysis of allele frequencies from the Geisinger MyCode dataset also indicates that selection is operating against eSDVs. Of the 14,633 MyCode variants which pass quality checks, 5,982 variants (40.9%) were only identified in a single patient sample. Due to the extreme rarity of these singleton variants, they have likely not been present in the population long enough for natural selection to act on them. Thus, we would expect deleterious variants such as stop gained variants to have a higher likelihood of being a singleton. We observe this enrichment for singletons among stop gained and splice region variants relative to synonymous variants (Figure 2b, left). We also find a higher rate of singletons within the lowest MaPSy splicing score percentiles, suggesting that purifying selection is acting against variants with the most severe splicing defects (Figure 2b, right). Among the assayed genes, *SCN3B*, *MLH1* and *BRCA1* had the highest concentration of eSDVs (Figure 2c). Defects in these genes are associated with the development of long QT syndrome, Lynch syndrome, and breast and ovarian cancers, respectively.

**Figure 2.**
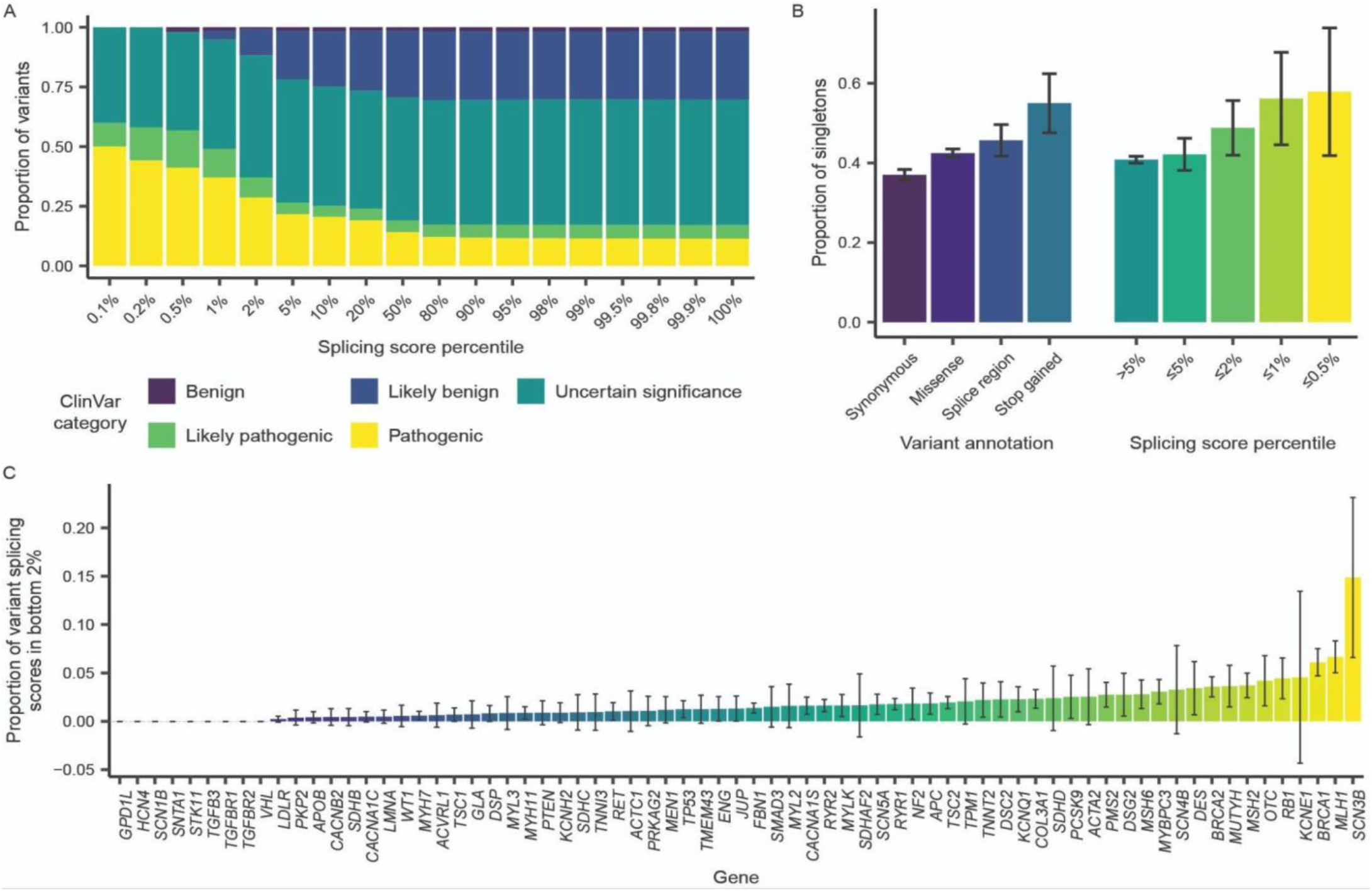
MaPSy splicing scores correlate with predicted variant pathogenicity and measures of selection. A) Tested variants that are present in ClinVar were stratified based on their MaPSy splicing scores. Each column contains the variants at or below the given splicing score percentile. B) The proportion of singleton variants from MyCode was calculated for variant sets based on their annotation as synonymous, missense, splice region, or stop gained variants by the Ensembl Variant Effect Predictor (left) or based on their MaPSy splicing score (right) (error bars represent ±SE). C) For the 71 clinically-actionable genes with variants that passed quality checks, the proportion of tested variants with splicing scores in the bottom 2% is plotted (error bars represent ±SE).

### Splice-disrupting variants are concentrated in hotspot exons

We have previously shown that SDVs distribute non-uniformly across genes, with certain exons being hotspots for splicing mutations^33^. This phenomenon is apparent in our splicing assay results. For instance, of the 15 assayed exons within *MLH1*, three exons had over 10% of tested variants categorized as eSDVs while five exons had no eSDVs whatsoever (Figure 3a). Analyzing multiple occurrences of a mutation within identical 5-mer motifs (e.g. all assayed instances of the TCGAC > TCAAC mutation) across different exons provides support for the importance of the exonic context in determining a mutation’s splicing outcome (Figure 3b). A given 5-mer mutation induces more severe splicing defects in exons with lower average splicing scores than the same mutation does in exons with higher mean splicing scores. To further assess the clustering of SDVs within particular exons, we compared the true distribution of exon mean splicing scores to the distribution computed after randomly permuting the associations between exons and variant splicing scores (100 trials). This analysis revealed an excess of exons with low mean splicing scores compared to the expectation if there was no relationship between the exon context and a variant’s effect (K-S test *P* = 7.96×10^-27^, Figure 3c). In order to assess the occurrence of this trend genome-wide, we performed an *in silico* analysis of every possible mutation in every exon of the CCDS annotation. Each mutation was scored for potential splice-disrupting effects with SpliceAI and the mean SpliceAI score for each exon was computed. After permuting the association between exon and a mutation’s SpliceAI score, the mean SpliceAI score was recalculated for each exon (100 trials). Comparing the true and permuted distributions again indicates the presence of hotspot exons with more predicted splice-disrupting mutations than the expectation (K-S test *P* = 0, Figure 3d).

**Figure 3.**
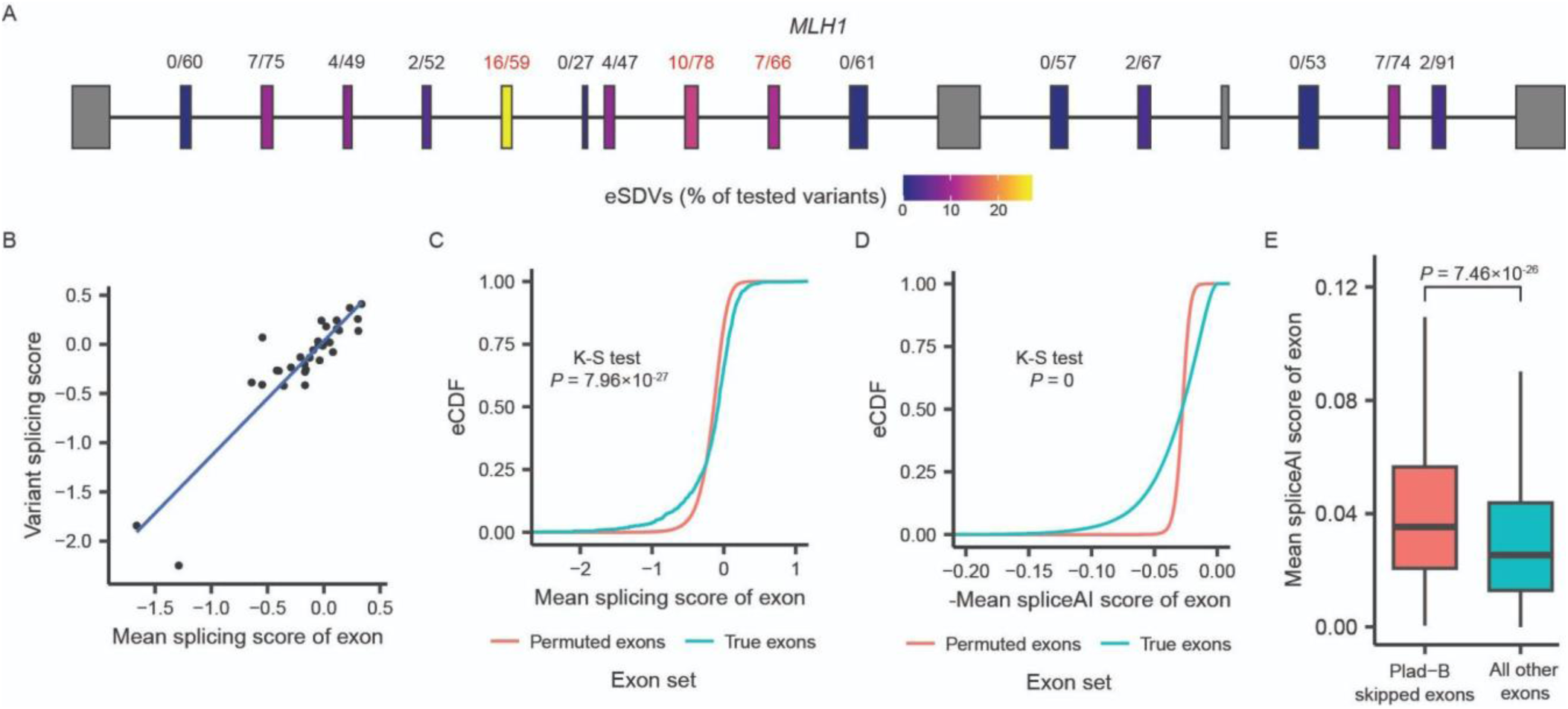
Hotspot exons are enriched for splice-disrupting variants and susceptible to inhibition by Pladienolide B. A) The distribution of extreme splice-disrupting variants (eSDVs, variants in bottom 2% of splicing scores) across assayed *MLH1* exons. eSDV and total tested variant count are given above each exon, and exons with >10% of tested variants being eSDVs are highlighted in red. B) For each of the 32 assayed G>A variants occurring in the context TCGAC > TCAAC, the variant splicing score is plotted against the mean splicing score of all variants in the same exon. C) The distribution of mean splicing scores for the assayed exons (true exons) compared to the mean splicing scores derived from 100 trials where the relationship between variant and exon was randomly permuted (permuted exons). D) The distribution of mean SpliceAI scores for the assayed exons (true exons) compared to the mean SpliceAI scores derived from 100 trials where the relationship between variant and exon was randomly permuted (permuted exons). E) The distribution of mean SpliceAI scores for exons which were skipped upon Pladienolide B (Plad B) treatment (change in exon inclusion ≤ −0.9) compared to all other exons.

Using a MaPSy splicing score threshold of −1.5 to define SDVs, we identify 83 of the 1074 assayed exons (7.7%) from the 71 actionable genes as being significantly enriched for SDVs (binomial test, FDR < 0.05). Having previously found that hotspot exons are sensitive to splice-altering drugs, we sought to identify potential hotspot exons through treatment of HEK293T cells with the general splicing inhibitor Pladienolide B (Plad B). Following RNA-seq of Plad B-treated and untreated samples, we performed differential splicing analysis using rMATS. Exons which had an inclusion change of ≤ −0.9 upon Plad B treatment were predicted by SpliceAI to be more susceptible to splicing mutations compared to exons that were not skipped in Plad B treated samples (Figure 3e).

### Antisense oligonucleotides reverse the exon skipping induced by a splicing inhibitor *in vivo*

In a previous study, we established that exonic features serve as robust predictors of the impact a variant will have on splicing^33^. The importance of the exonic context suggests that most splicing mutations will be discovered in the minority of exons which are most susceptible to splicing disruption. This also suggests a treatment that improves context could revert any mutation in that exon. It has been shown that an exon’s inclusion is influenced by the strength of its flanking exons^33^. As exon skipping is a competition between splice sites, we hypothesize that ASOs targeting the flanking splice sites (i.e. the sites involved in exon skipping) could act to promote inclusion. This increased inclusion could rescue splicing defects caused by any mutations in hotspot exons. To test this hypothesis, we identified 5 exons which were susceptible to disruption by Plad B and treated them with ASOs targeted to the upstream 5’ splice site (5’ss) or downstream 3’ splice site (3’ss) (Figure 4a). The rate of skipping upon Plad B treatment was between 13-96% for these exons, and co-treatment with one or both of the flanking splice site ASOs reverted between 27-86% of the Plad B-induced skipping (Figure 4b-f). Treatment with a single ASO tended to result in similar levels of splicing reversion as simultaneous treatment with both 5’ss and 3’ss ASOs.

**Figure 4.**
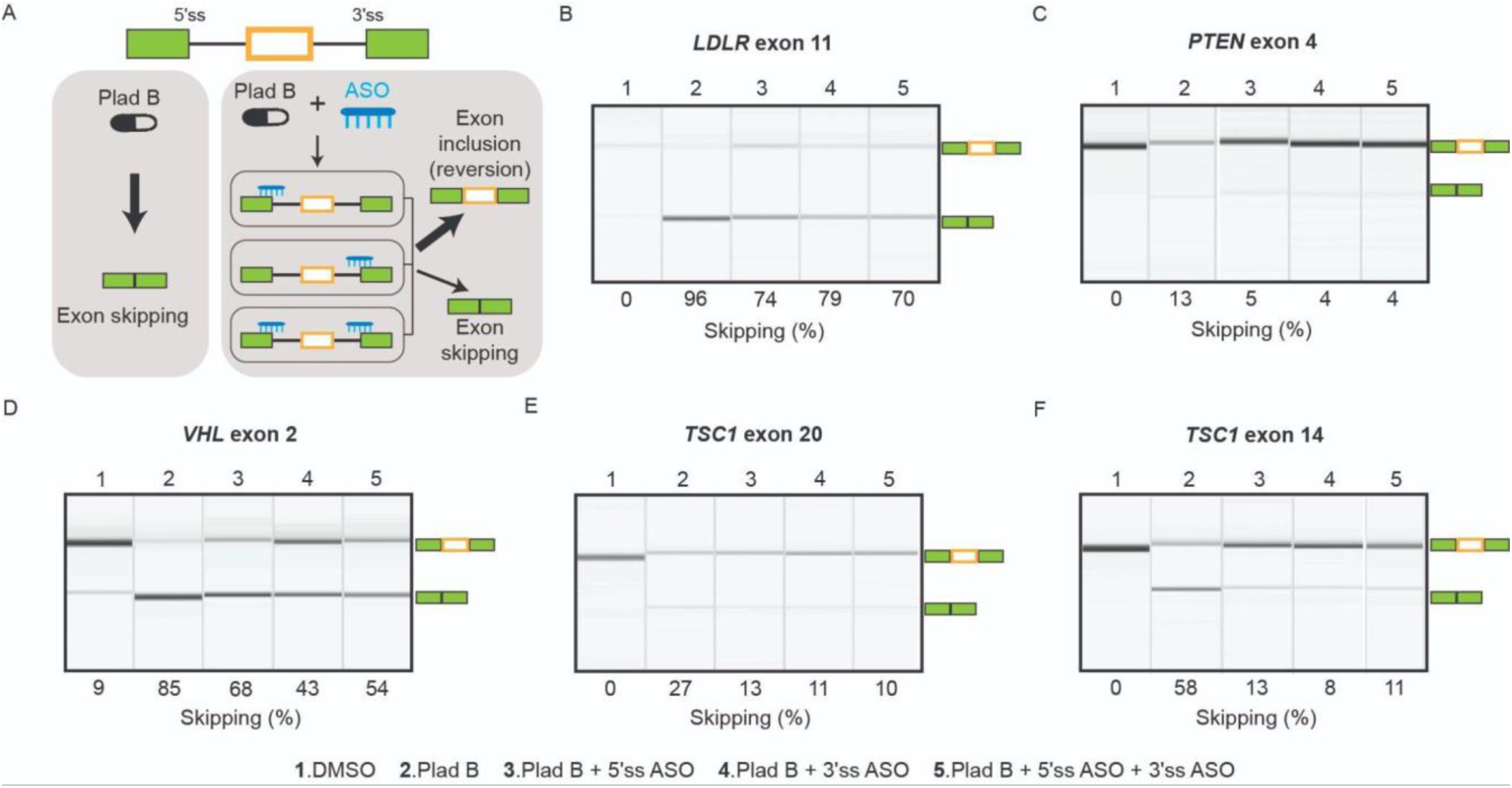
Antisense oligonucleotides reverse the exon skipping induced by a splicing inhibitor *in vivo*. A) Five exons (orange box) in actionable genes sensitive to Pladienolide B (Plad B) were selected for antisense oligonucleotide analysis as depicted. Endogenous genes exposed to Plad B were treated with ASO (blue) targeting one or both of upstream 5’ splice site (5’ss) and downstream 3’ splice site (3’ss) in flanking exons (solid green). B-F) RNA extracted from treated HEK293T cells was analyzed by PCR. Exon inclusion and skipping bands (cartoon, right side) were quantified (below each gel). Lane number indicates treatment (lane treatment below panel).

### Antisense oligonucleotides revert the effects of splice-disrupting variants

Having observed the ability of ASOs to counteract drug-induced splicing effects, we sought to test whether they could rescue exon skipping caused by SDVs discovered through MaPSy. We selected *TSC1* exon 14 and *MLH1* exon 9 as test hotspot exons and constructed splicing reporters containing the exon of interest in either wildtype or variant forms flanked by the endogenous upstream and downstream exonic and intronic sequence context (Figure 5a). Following transfection in HEK293T, the splicing products were analyzed by RT-PCR using primers in the upstream exon and polyA sequences (Figure 5b-c). Variants with <10% exon skipping (*TSC1* variant 4 and *MLH1* variants 4-5) were excluded from subsequent ASO rescue tests. For the 3 remaining variants in each exon, variant minigenes were co-transfected with ASOs targeting the upstream 5’ss and downstream 3’ss (Figure 5d-e). For the *TSC1* variants, ASOs were also tested against the branchpoint (BP) of the downstream intron (Figure 5d, BPv1/BPv2). ASO treatment reverted 89-100% of the exon skipping induced by the variants in *TSC1* and 72-100% of the exon skipping caused by the *MLH1* variants. While the ASOs targeted to the branchpoint of the downstream intron performed with similar efficacy as those targeted to flanking splice sites, the design of branchpoint ASOs is limited by the fact that many introns lack a published branchpoint annotation. Overall, these findings confirm the ability of single ASOs to revert the skipping caused by splice-disrupting variants.

**Figure 5.**
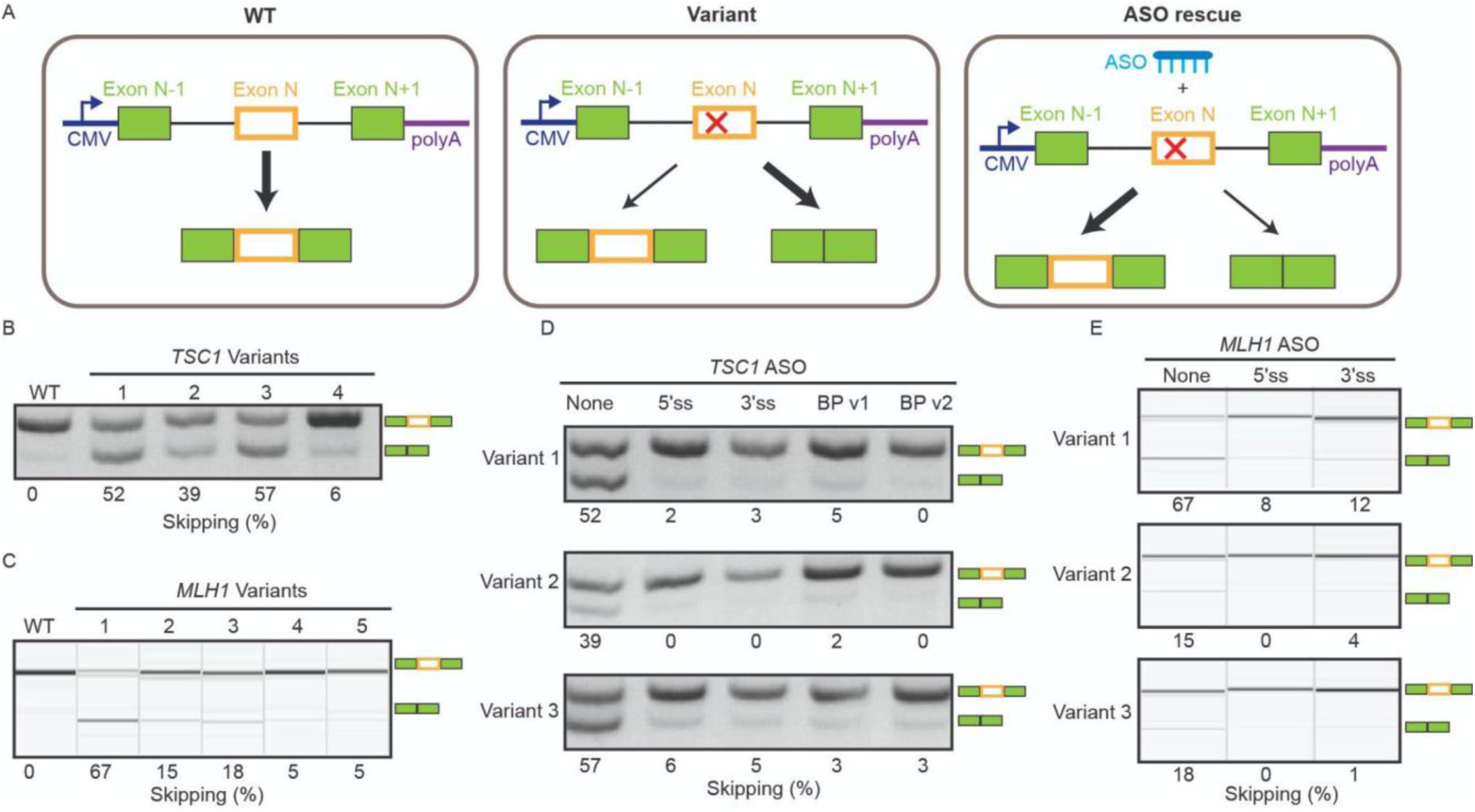
Antisense oligonucleotides revert the effects of splice-disrupting variants. A) For *TSC1* exon 14 and *MLH1* exon 9, variants discovered to alter splicing in MaPSy (Figure 1) were engineered into minigene reporters containing the endogenous sequence context between the upstream and downstream flanking exons and B, C) validated by PCR. D) ASO complementary to the following targets: upstream 5’ss, downstream 3’ss, or downstream branchpoint (BP v1, BP v2) were tested for 3 variants in *TSC1* exon 14. E) Upstream 5’ss and downstream 3’ss ASOs were tested for 3 variants in *MLH1* exon 9.

## DISCUSSION

In this work, we assayed 31,112 variants from ClinVar and Geisinger MyCode that overlap internal exons of 71 clinically actionable genes and identified 1,733 SDVs. The reporter assay allows for the comparison of the relative representation of the mutant to the wildtype in the spliced output versus input. Furthermore, we identified 83 hotspot exons from the 71 clinically actionable genes in this study. Intriguingly, our prior research revealed that exon level features contribute more to the prediction of splicing mutations than the features of the mutation itself (e.g. base change, disrupted motif)^33^. The findings presented here suggest these exon features create tenuous splicing substrates where numerous mutations are capable of disabling the recognition event. An important consequence of this model is that a treatment that strengthens the splicing of an exon could revert any mutation that disrupts its splicing. As exon skipping is a competition between splice sites, we reasoned that inhibiting the splice sites used in the skipping event could theoretically increase the inclusion of the affected exon. We demonstrate, for all 5 exons tested, that ASOs targeted against flanking exons restored the exon inclusion defects caused by splicing variants. For hotspot exons, mutations that disrupt splicing could also be phenotypically reverted by a similar ASO strategy.

To our knowledge, this represents the first principled approach to increasing the splicing of a particular substrate. The proposed ASO strategy appears to work on all the alleles in a particular exon. For example, our findings indicate that a single ASO targeting the flanking exon of *TSC1* exon 14 can effectively reverse exon skipping induced by any of the three mutations that disrupt splicing. This phenomenon is likely attributed to the ASO’s capacity to attenuate competing splice sites. Irrespective of the specific target locus of ASOs, such as the upstream 5’ splice site or the downstream 3’ splice site, as long as the ASOs can suppress the competing splice site, it proves effective. It’s not surprising that two ASOs targeting the branchpoint region of the downstream intron also rescue the exon skipping.

In the context of rare heritable diseases, where traditional drug development may not be economically viable due to the small patient population, ASOs hold considerable potential. For instance, in Duchenne muscular dystrophy (DMD), ASOs are used to induce exon skipping, restoring the reading frame and enabling dystrophin expression. Several ASOs, such as Casimersen, Eteplirsen, Golodirsen, and Viltolarsen, which target exons 45, 51, and 53 of the *DMD* gene, have been approved by the FDA^22,34–36^. In the case of spinal muscular atrophy (SMA), the FDA has approved Nusinersen, an ASO that binds specifically to an intronic splicing silencer within intron 7, thereby preventing the binding of splicing repressors hnRNP A1/A2 and promoting exon 7 inclusion in the final *SMN2* mRNA transcript^37–39^. Initial efforts to enhance the inclusion of exon 7 in *SMN2* mRNA involved an ASO designed to block the 3′ splice site of exon 8^18^, similar to the approach used in this study. Although this strategy ultimately did not result in the development of a drug, it successfully promoted exon 7 inclusion. Our work focused on hotspot exons, showing that a single ASO can address multiple splicing defects within the same exon, providing a versatile, one-size-fits-many therapeutic solution. By enabling the correction of splicing for various mutations, a single ASO can potentially treat a broader range of patients with the same underlying disorder.

## METHODS AND MATERIALS

### Selection criteria for variants in MaPSy library

An initial list of 76 clinically actionable genes from Dewey et al^29^ was considered. For each gene, a single canonical transcript was retained based on APPRIS principal transcript annotations^40^ from all GENCODE v32 basic transcripts^41^. MyCode variants identified in 140K patient exomes and ClinVar variants (ftp://ftp.ncbi.nlm.nih.gov/pub/clinvar/vcf_GRCh38/clinvar_20200120.vcf.gz, as of 2020-01-20)^42^ were overlapped with internal exons of each gene’s canonical transcript. For short exons (10 ≤ length ≤ 120 bp), we retained 15,349 exonic variants spanning the full width of each exon. For long exons (length > 120 bp), we retained 21,122 exonic variants within the first 90 bp next to the 3’ss of each exon. In total, we retained 19,702 variants in the MyCode 140K exome dataset and 25,761 variants in ClinVar overlapping 71 clinically actionable genes (36,471 variants in total). These variants overlap 71 out of the initial set of 76 genes. (*PLN*, *CAV3*, *KCNE2*, and *KCNJ2* did not have internal exons, and *KCNE3* did not overlap any variants.)

### MaPSy library transfection and sequencing

The oligo library design for the selected ClinVar variants and the construction of MaPSy minigenes, as described by Rong et al.^43^, were implemented. The resulting MaPSy minigene input libraries were transfected into HEK293T cells obtained from the American Type Culture Collection in three cell culture replicates using Lipofectamine 3000 (Invitrogen) in a 6-well plate. 24 hours after transfection, RAN was extracted by RNeasy Mini Kit (Qiagen), followed by DNase treatment (Invitrogen). Random 9-mers were used to generate cDNA with SuperScript IV Reverse Transcriptase (Invitrogen) followed by PCR (GoTaq, Promega), resulting in output libraries of spliced species. Input and output libraries were sequenced using Illumina HiSeq 2×150bp.

### Alignment and QC of MaPSy sequencing data

STAR^44^ was used to perform end-to-end unspliced alignment of paired-end reads against custom reference genomes with either the expected input minigene sequences or the expected output exon inclusion sequences with up to 5 mismatches and no indels allowed, using the parameters: -- alignIntronMax 1 --outFilterMismatchNmax 5 --scoreDelOpen -1000 --scoreInsOpen -1000 -- alignEndsType EndToEnd --outFilterMultimapNmax 1. SAMtools idxstats^45^ was used to count the number of reads corresponding to each input and output species in each replicate sequencing library. The resulting read counts were then joined to a table of variant-exon pairs.

We performed quality control (QC) of variant-exon pairs based on the following filters: a minimum input wildtype and mutant read counts of 100 in all three replicates, a minimum output wildtype read count of 100 in all three replicates, and at least one output mutant read in one of the three replicates. After QC, we retained 17,337 variants in MyCode and 22,686 variants in ClinVar (32,112 variants in total) for analysis.

### Analysis of read count data

The MaPSy splicing score is defined as the log fold change of mt versus wt read counts in the output library, normalized by read counts in the input library, or 𝑙𝑜𝑔_2_((𝑚𝑡_𝑜_/𝑤𝑡_𝑜_)/(𝑚𝑡_𝑖_/𝑤𝑡_𝑖_)). To estimate the MaPSy splicing score using three replicates, log fold change (FC) estimates from mpralm^46^, a variation of the limma-voom framework for allelic differential expression in massively parallel reporter assays^47,48^, were utilized. The mpralm log FC estimates and the mpralm moderated t-test p-values with FDR correction for multiple tests ^49^ were employed. Significant SDVs were identified at FDR < 0.05 and |FC| > 1.5. Variant-exon pairs were then ranked by MaPSy splicing score to identify SDVs at varying splicing effect percentile thresholds, with a particular focus on variants below the 2%-tile.

### Pladienolide B treatment

Pladienolide B (Plad B) was purchased from Cayman Chemical and dissolved in DMSO. HEK293T cells were treated with either DMSO or 250 nM Plad B for 24 hours. Total RNA was extracted using the RNeasy Mini Kit (Qiagen), and submitted to Genewiz for next-generation sequencing.

To test for exon skipping following Plad B treatment, 1 μg of total RNA was reverse-transcribed using SuperScript IV Reverse Transcriptase (Invitrogen), followed by PCR amplification with GoTaq (Promega). The primers are listed in Table 1.

**Table 1.**
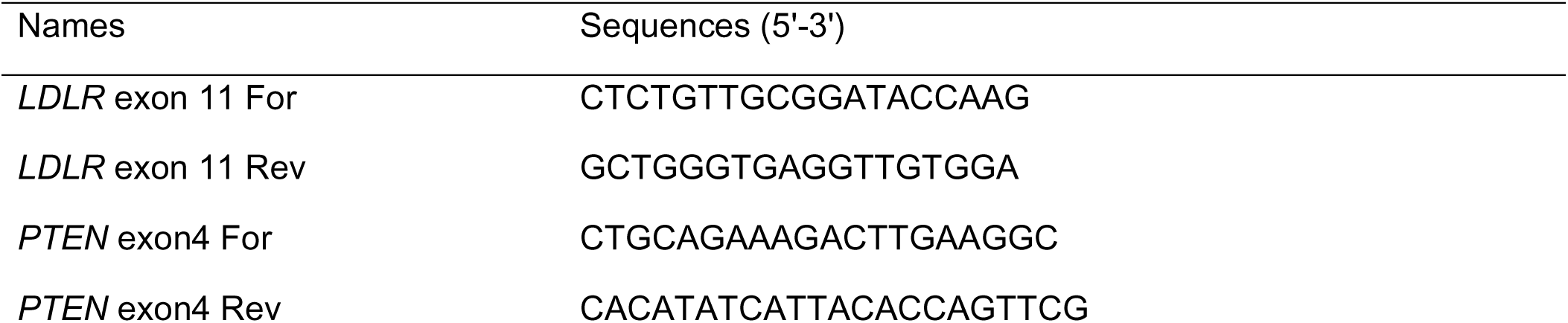
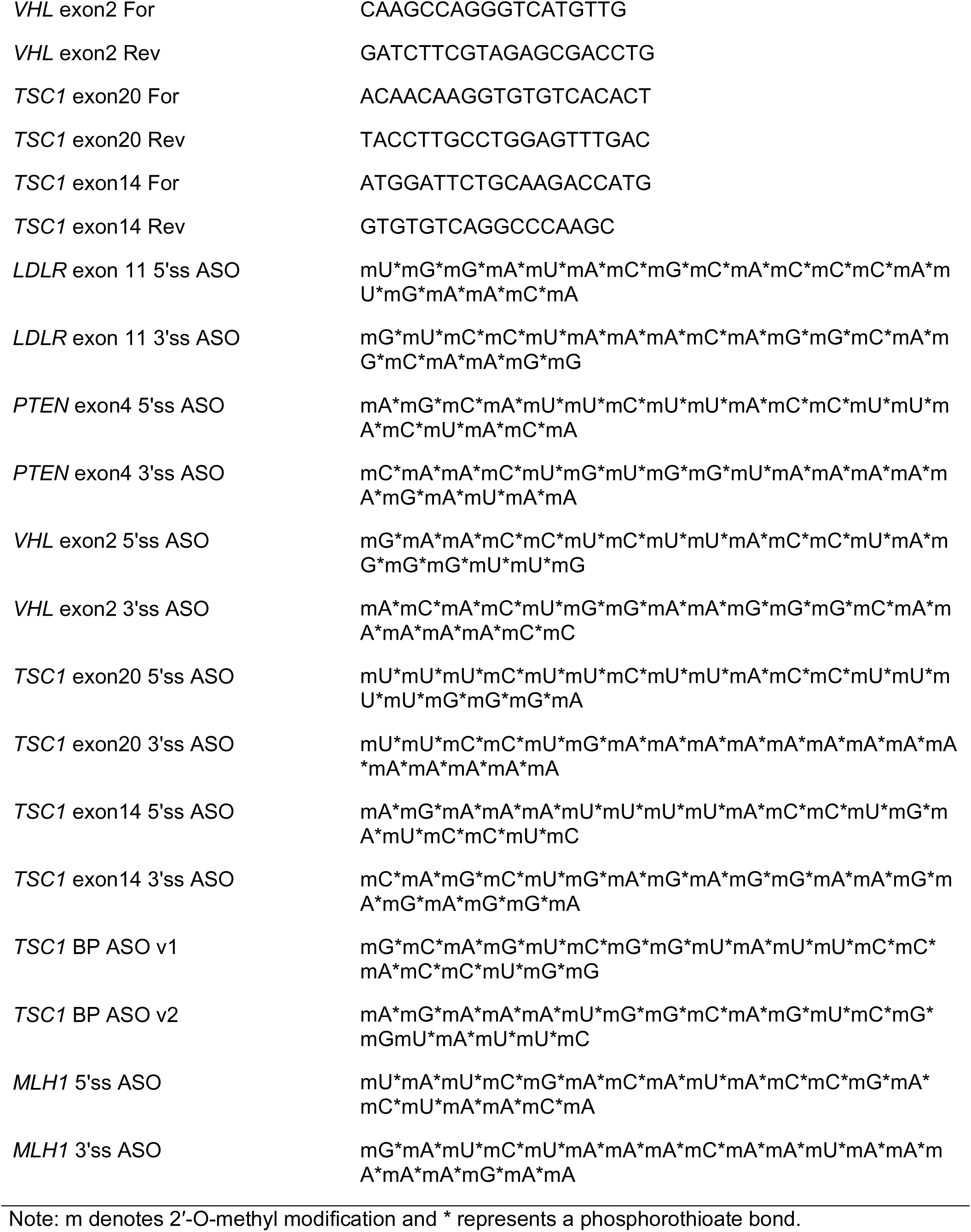
Sequences of primers and antisense oligonucleotides used in this study.

Antisense oligonucleotides (ASOs) were designed to target either the upstream 5’ splice site or the downstream 3’ splice site of the skipped exon. ASOs were synthesized by GenScript (Table 1) with phosphorothioate backbones and 2’-O-methyl modifications on every base. HEK293T cells were transfected with 300 nM of ASO using Lipofectamine 3000 (Invitrogen). Six hours after ASO transfection, the medium was changed to one containing either DMSO or 250 nM Plad B. Cells were harvested for RNA extraction 24 hours after Plad B treatment.

### *TSC1* and *MLH1* minigene construction and antisense oligonucleotides experiments

The gene fragments for *TSC1* and *MLH1* (exons along with their flanking introns) were synthesized by Twist Bioscience. The CMV promoter and bGH polyA sequences were then assembled using overlap PCR. For the *TSC1* minigene construction, the entire lengths of the relatively small upstream and downstream introns of exon 14 (465 bp and 591 bp, respectively) were included in the minigene reporters. For the *MLH1* minigene construction, the upstream and downstream introns of exon 9 are much larger (2332 bp and 2961 bp, respectively). Therefore, only 100 bp from the 5’ end and 200 bp from the 3’ end of these introns were incorporated into the minigene reporters. In both cases, the complete upstream and downstream exons were included.

HEK293T cells were seeded in a 6-well plate the day before transfection to achieve ∼70-80% confluence at the time of transfection. Transfection was carried out with 300 nM of ASO and 0.5 μg of minigene per well using Lipofectamine 3000 (Invitrogen), following the manufacturer’s instructions for forward transfection. Cells were harvested 24 hours after transfection for total RNA extraction. The ASOs are listed in Table 1.

## AUTHOR CONTRIBUTIONS

C.D. and C.R.N. performed experiments. S.R., L.B. and C.D. performed data analysis. J.M.S and N.T.S collected and systematically organized patient sequencing data for analysis. C.D., L.B., and W.G.F. wrote the manuscript. W.G.F. conceived and supervised the study.

## ACKNOWLEDGEMENT

We would like to acknowledge the participants of the MyCode Community Initiative for the use of their health and genomic information, without whom this study would not be possible. The patient enrollment and exome sequencing were funded by the Regeneron Genetics Center. Data for this project was made possible by the Geisinger-Regeneron DiscovEHR Collaboration.

---

**Supplementary Figure 1.**
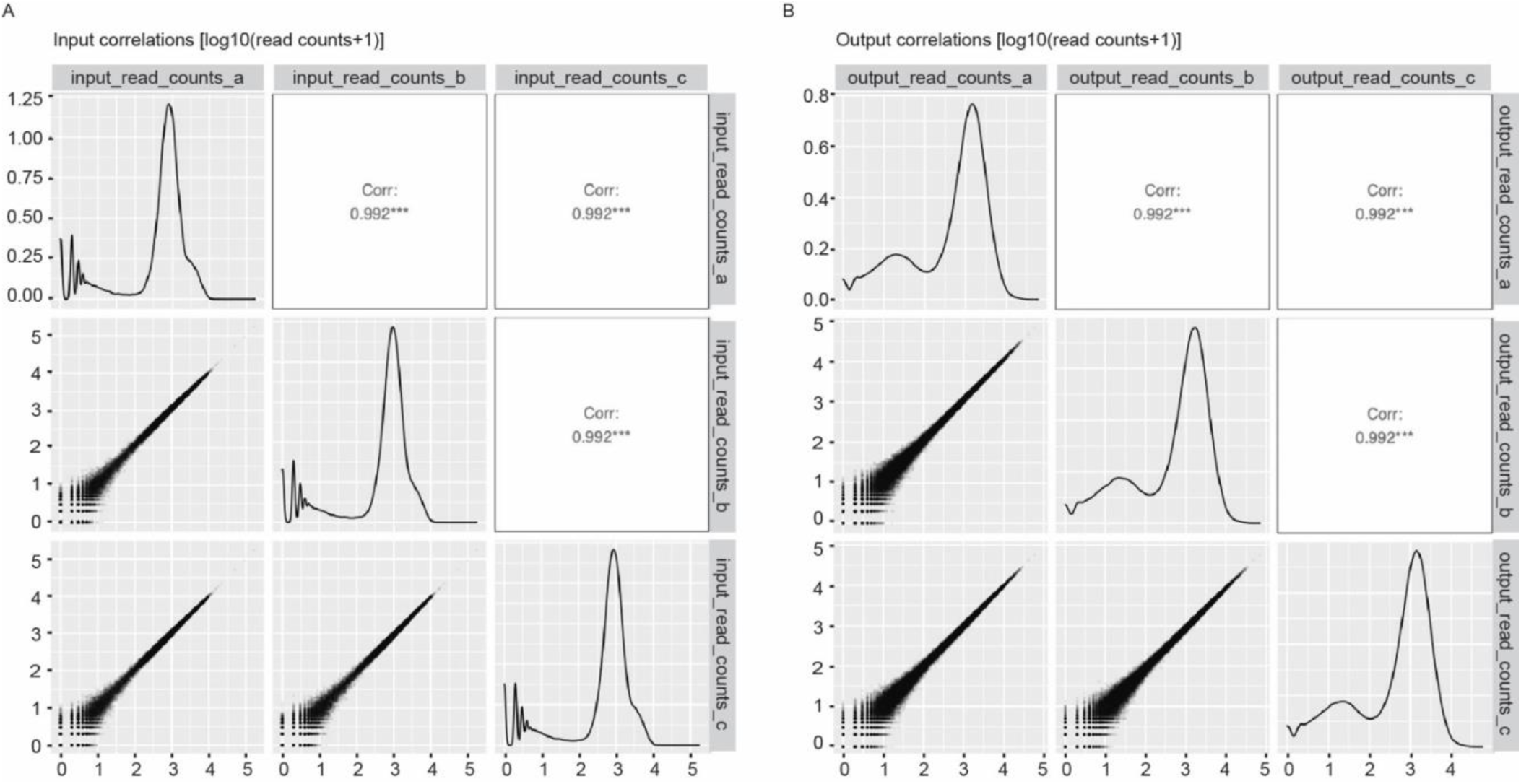
High correlation of sequencing read counts (log10(read counts + 1) among three biological replicates. A) Input MaPSy sequencing replicates. B) Output MaPSy sequencing replicates.

**Supplementary Figure 2.**
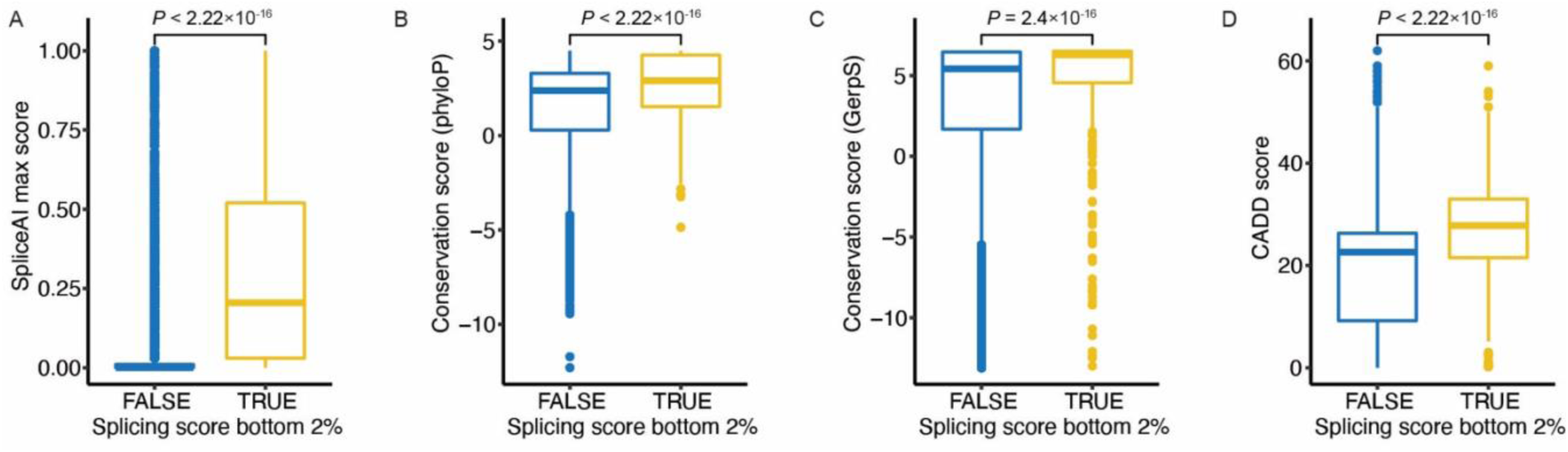
Variants in bottom 2% of splicing scores are significantly enriched in predicted splicing effects and evolutionary conservation. Comparing distributions for variants below versus above 2%-tile for: A) SpliceAI max score, B) mammalian phyloP score, C) GerpS score, and D) CADD score. *P*-values from Kruskal-Wallis test.

